# Venom diversity in *Naja mossambica*: Insights from proteomic and immunochemical analyses reveal intraspecific differences

**DOI:** 10.1101/2023.10.27.564320

**Authors:** Konrad K. Hus, Justyna Buczkowicz, Monika Pietrowska, Vladimír Petrilla, Monika Petrillová, Jaroslav Legáth, Thea Litschka-Koen, Aleksandra Bocian

## Abstract

**Background:** Intraspecific variations in snake venom composition have been extensively documented, contributing to the diverse clinical effects observed in envenomed patients. Understanding these variations is essential for developing effective snakebite management strategies and targeted antivenom therapies. This study was prompted by the observations made by clinicians, who have noted significant variations in clinical outcomes among patients bitten by *Naja mossambica* in different regions of Africa, which links to the phenomenon of intra-species venom variability. We aimed to comprehensively investigate venoms from three distinct populations of *N. mossambica* from Eswatini, Limpopo, and KwaZulu-Natal regions in Africa in terms of their protein composition and reactivity with three commercial antivenoms (SAIMR polyvalent, EchiTAb+ICP, and Antivipmyn Africa).

**Methodology/Principal Findings:** In contrast to previous reports, we discovered an unexpectedly high concentration of neurotoxic proteins in *N. mossambica* venoms (approximately 15%). The Eswatini population of Mozambique spitting cobra exhibited an increased abundance and diversity of neurotoxic proteins, including neurotoxic 3FTxs, kunitz-type inhibitors, vespryns, and mamba intestinal toxin 1.

Immunochemical assessments of venom-antivenom reactivity unveiled differences, primarily related to low-abundance proteins. Notably, the reactivity of EchiTAb+ICP antivenom surpassed that of the widely used SAIMR polyvalent in serial dilution ELISA assays.

**Conclusions/Significance:** Our findings reveal a substantial presence of neurotoxic proteins in *N. mossambica* venoms, challenging previous understandings of their composition. Additionally, the detection of numerous peptides aligning to uncharacterized proteins or proteins with unknown functions underscores a critical issue with existing venom protein databases, emphasizing the substantial gaps in our knowledge of snake venom protein components. This underscores the need for enhanced research in this domain. Significantly, our research highlights the superior reactivity of EchiTAb+ICP antivenom compared to SAIMR polyvalent, providing another compelling argument for its potential as an alternative to the commonly used SAIMR antivenom.

**Author Summary:** Snakebite envenoming is a pervasive global health concern, posing substantial risks, particularly in less developed regions. The intricate variations in venom composition within a single species have been well-documented, contributing significantly to the varied clinical effects experienced by envenomed patients. It is imperative to unravel these variations, as they are pivotal in the formulation of effective snakebite management strategies and the development of targeted antivenom therapies.

In this study, our focus rested on the venom of the *Naja mossambica* species, dwelling in diverse African regions. Our objective was to delve into the toxin composition of these venoms and understand how these toxins interact with commercially available antivenoms. This exploration was aimed to uncover which toxins, despite antivenom application, evade neutralization. This information becomes a cornerstone in the design of more potent and efficacious antivenoms, contributing to a nuanced approach in combating the complex landscape of snakebite envenoming.

## 1. Introduction

Snakebite envenoming is a significant global health issue, causing substantial morbidity and mortality, particularly in developing regions. Annually, an estimated 81,000 to 138,000 people die from snakebites, with approximately four times as many suffering severe injuries [1–2]. Sub-Saharan Africa alone witnesses an annual occurrence of around half a million envenoming cases, which represent over 20% of the reported snakebite envenoming incidents worldwide. This region’s high incidence contributes to an estimated 32,000 fatalities, although these figures likely underestimate the true impact due to underreporting, especially in remote areas with limited access to medical care [3–4].

Venom diversity plays a crucial role in determining the clinical manifestations of snakebite envenoming. Variations in venom composition have been extensively documented not only between different snake species but also, more recently, within a single species [5–11]. Hence, intraspecific differences in venom composition have emerged as an important area of study, highlighting the complex nature of venom. Factors such as geographic location, diet, genetic variation, and evolutionary adaptations can contribute to these variations [2,11–12]. Gaining a comprehensive understanding of these intraspecific differences is essential for developing effective snakebite management strategies and targeted antivenom therapies, as they directly impact the diverse clinical effects observed in envenomed patients [2,12].

*Naja mossambica*, also known as the Mozambique spitting cobra, serves as an interesting model for investigating such intraspecific variations in venom composition. This species primarily inhabits the lowland areas of sub-Saharan Africa, with a distribution ranging from Tanzania in the north to Angola and Namibia in the west, and the KwaZulu-Natal province in the south [13]. The venom of the Mozambique spitting cobra is recognized for its primarily cytotoxic effects, characterized by the prevalence of cytotoxic three-finger toxins, constituting approximately 70-80% of all venom proteins. Additionally, phospholipases A_2_, the second most abundant group of proteins, account for approximately to about 10-30% of all toxins in *N. mossambica* venom [14–16]. Currently, the treatment of *Naja mossambica* snakebites in Sub-Saharan Africa relies solely on the SAIMR polyvalent antivenom, posing significant challenges and risks for the local population. However, recent preclinical findings by Menzies et al. suggest that other antivenoms also hold the potential for neutralizing the venom of this snake, offering a chance to improve the situation and reduce reliance on a single antivenom [17]. However, further data in this area are needed to better understand the effectiveness of alternative antivenoms.

In this study, our objective is to comprehensively characterize the intraspecific variations in venom composition among three distinct populations of *Naja mossambica* originating from the regions of Eswatini, Limpopo, and KwaZulu-Natal, and assess their impact on the interactions with three commercially available antivenoms (SAIMR polyvalent, EchiTAb+ICP, Antivipmyn Africa).

## 2. Materials and methods

### 2.1. Venom and antivenom collection

Venom samples from the species *Naja mossambica* were collected following approval by The Ethics Committee at the UVMP in Košice, permission No. EkvP/2023-09. The samples were taken at the breeding facilities VIPERAFARM with registration number CHNZ-01-TT, Tropický svet with registration number CHEZ-TT-01, and UVMP with registration number SK NZ 0010/2022 by a specialist certified by SNTC/FAGASD. The examined individuals kept in these breeding facilities originated from specific locations, namely Limpopo (country: South Africa; province: Limpopo; city: Ellisras), KwaZulu-Natal (country: South Africa; province: KwaZulu-Natal; city: Pietermaritzburg), and Eswatini (Country: Eswatini; Province: Manzini; City: Manzini). Individuals in the breeding facilities were captured using standard methods with protective handling equipment. A total of 15 individuals of the species *Naja mossambica* were available for venom collection, comprising 5 individuals from the Limpopo region, 6 individuals from the KwaZulu-Natal region, and 4 individuals from the Eswatini region. Snake venoms were extracted directly into glass collection containers and subsequently placed in microcentrifuge tubes (1.5 ml). Venom extraction was conducted from adult and subadult individuals of both sexes every two weeks over the course of three months. Samples collected at the same time from the snakes that originated from the same region were combined, resulting in six venom samples for each region. During transportation, venom samples were stored at −20 °C and then stored in a deep freezer box at −80 °C.

EchiTAb+ ICP (commonly known as PanAfrican Antivenom) (LOT:5851216 DEPA-C; Expiry date: 12/21) and Antivipmyn Africa (LOT: B-9B-14; Expiry date: 02/12) antivenoms were provided by the Eswatini Antivenom Foundation for this study, whereas SAIMR Polyvalent Snake Antivenom (LOT:BM00746; Expiry date: 02/24) was purchased directly from the manufacturer (South African Vaccine Producers).

Protein concentration in venom and antivenom samples was measured using Pierce™ BCA Protein Assay Kit (Thermo Fisher Scientific, Waltham, MA, USA) according to the manufacturer’s instructions. The respective measured concentration values were as follows: Antivipmyn Africa – 8.67 µg/µL; EchiTAb+ ICP – 181.88 µg/µL; SAIMR Polyvalent – 206.33 µg/µL.

### 2.2. Whole venom shotgun LC-MS/MS analysis

18 venom samples (six from each region) containing 50 µg of proteins were desiccated using a SpeedVac Vacuum Concentrator (Thermo Fisher Scientific, Waltham, MA, USA). The proteins were reduced utilizing a 5 mM solution of tris-2-carboxyethyl-phosphine (TCEP) (Merck KGaA, Darmstadt, Germany) at a temperature of 60 °C for 1 hour. Subsequently, the reduced cysteine residues were blocked employing a 10 mM solution of S-methyl methanethiosulfonate (MMTS) (Merck KGaA, Darmstadt, Germany) at room temperature for 10 minutes. Trypsin (Promega, Madison, WI, USA) was added at a 1:25 (v/v) ratio, and the samples were incubated overnight at 37 °C to facilitate enzymatic digestion. To inactivate trypsin, 0.1% trifluoroacetic acid (TFA) (Merck KGaA, Darmstadt, Germany) was added.

Peptide identification and absolute quantitation were performed using nanoACQUITY UPLC system (Waters Corporation, Milford, MA, USA) coupled to Q Exactive mass spectrometer (Thermo Fisher Scientific, Waltham, MA, USA). The samples were loaded onto the nanoACQUITY UPLC Trapping Column (Waters Corporation, Milford, MA, USA) using a mobile phase consisting of water containing 0.1% formic acid (Merck KGaA, Darmstadt, Germany). Subsequently, the samples were separated on the nanoACQUITY UPLC BEH C18 Column (Waters Corporation, Milford, MA, USA, 75 µm inner diameter; 250-mm long) using an acetonitrile gradient (5–35% acetonitrile for 160 min, Merck KGaA, Darmstadt, Germany) in the presence of 0.1% formic acid at a flow rate of 250 nl/min. High-energy collisional dissociation (HCD) fragmentation was employed. Up to 12 MS/MS events were considered per MS scan, and the spectrometer resolution was set to 17,500. Whole venom shotgun LC-MS/MS experiment was conducted at the Mass Spectrometry Laboratory, Institute of Biochemistry and Biophysics, Polish Academy of Sciences, Warsaw, Poland.

The acquired MS/MS raw data files were analyzed using MaxQuant software (ver. 2.2.0.0, Jürgen Cox, Max Planck Institute of Biochemistry, Martinsried, Germany) [18]. Protein identification was performed against the UniProtKB Serpentes database (release 12/2022) using the Andromeda engine. Methylthio (C) (+45.9877 Da) was set as a fixed modification, while oxidation (M) (+15.9949 Da) and acetyl (protein N-term) (+42.0105 Da) were considered variable modifications. Mass tolerance was set to 20 ppm for the initial MS search, 4.5 ppm for the main MS search, and 20 ppm for MS/MS fragment ions. Trypsin with full specificity and a maximum of two missed cleavages was used as the enzyme. The PSM and protein false discovery rate (FDR) were set to 1%. Further data analysis was performed with Perseus software (ver. 1.6.7.0) [19]. Hits identified only by site, those found in decoy or contaminant lists, were subsequently filtered out. Moreover, only proteins identified in at least three venom samples of the same region were further subjected to quantitative and qualitative proteome analysis. To account for the limitations of database-based identification methods which are more apparent when studying organisms with substantial gaps in the target database, we decided to deviate from the typical proteomic principle of considering identifications based on a minimum of two peptides. Instead, we also included proteins identified based on a single peptide, and assigned them to certain protein families, which were a subject of further quantitative and qualitative analysis. Although this approach influenced the obtained results, we deemed it more appropriate than rejecting numerous peptides that were solely assigned to specific proteins. The presence of such peptides may be attributed to being fragments of proteins with unknown yet sequences, which are absent in the target database but share identical peptides with other toxins. The intensity-based absolute quantification (iBAQ) values of razor and unique peptides were used to calculate the amount of each specific protein in the sample. Non-zero iBAQ values from three to six samples were averaged and assigned to the corresponding proteins. The proteins were then categorized into different protein groups/families, and the percentage of each protein group was determined by dividing the summed iBAQ values of proteins in the group by the summed iBAQ values of all quantified proteins identified in the sample. Based on these percentage data, pie charts illustrating the protein family distribution within each venom were generated.

To find the proteins whose amount differed between samples from different regions, statistical analysis was performed in Perseus software. The data was log(2)-transformed and missing values were imputed from normal distribution. Principal component analysis (PCA) was performed in Persues to test homogeneity of the samples within the groups. Multiple sample t-test with permutation-based testing correction (250 randomizations) was applied to find differences in protein abundance between samples. A false discovery rate threshold of 0.05 was employed. Subsequently, hierarchical clustering of proteins that passed the test was performed using the software. The clustering was based on logarithmized iBAQ values after z-score normalization, utilizing Euclidean distances with default parameters. The resulting clustering analysis included four clusters for the row tree, and for each cluster annotation enrichment analysis were performed with Benjamini-Hochberg FDR set at 0.04. To visualize the number of identified peptides in each sample, Venn diagram was prepared in InteractiVenn software [20].

### 2.3. Two-dimensional electrophoresis and 2D Western Blot

Six venom samples from a given region were pooled with equal mass ratios, resulting in three pooled region-specific samples that were used in further analyses.

For isoelectrofocusing, the samples were prepared by suspending 300 μg of proteins in 300 μl of thiourea rehydration solution comprising of 7 M urea, 2 M thiourea, 2% (v/v) Nonidet P-40, 0.5% (v/v) IPG buffer (pH range 3-10), 0.002% bromophenol blue, and 18 mM DTT (samples for silver staining contained 600 μg of proteins). The rehydration and isoelectric focusing steps were performed using 17 cm, pH 3-10 ReadyStrip IPG Strips (Bio-Rad, Hercules, CA, USA) at 50 μA per strip at 20 °C, according the following program: 12 h of active rehydration at 50 V and focusing: rapid ramp to 250 V – 30 min, linear gradient to 10,000 V – 3 h, and rapid ramp until 40,000 VHr, using PROTEAN IEF Cell device (Bio-Rad). Prior to SDS-PAGE, the strips were equilibrated for 10 min equilibration buffer solutions (6 M urea, 75 mM Tris-HCl pH 8.9, 30% (v/v) glycerol, 2% SDS, 0.002% bromophenol blue) first containing 65 mM DTT and second 216 mM iodoacetamide instead of DTT. In order to fit the strips into the electrophoresis apparatus, an approximately 2.5 cm fragment was cut from the anode part of the strip, hence pH values on the gels’ and membranes’ images start at approximately pH 4.5. The strips were immobilized on top of the gel with 1% agarose containing 0.002% bromophenol blue. The BlueEasy Prestained Protein Marker was utilized as a mass standard. SDS-PAGE was performed on 15% polyacrylamide gels (16 x 17.5 cm) with TGS buffer, using omniPAGE WAVE Maxi system (Cleaver Scientific, Warwickshire, UK). Protein spots were visualized by Coomassie Brilliant Blue G-250 or by silver staining according to protocol presented by Shevchenko [21].

For Western Blot analysis, wet electrotransfer of proteins from the gels to a 0.2 μm or 0.45 μm nitrocellulose membranes was performed using TG buffer containing 5% (v/v) of methanol. Subsequently, the membranes were incubated in a blocking solution consisting of 5% (w/v) skim milk in PBS/T (0.05% (v/v) Tween-20) for 2 hours. Following the blocking step, the membranes were exposed to primary antibodies (individual antivenoms) suspended in the blocking solution at a final mass of 2.1 mg, and the incubation was carried out overnight. Afterward, membranes were incubated with HRP-labeled anti-horse secondary antibodies diluted in PBS/T at a ratio of 1:1250, for 3h. Signal detection was carried out using a DAB substrate solution containing 0.5 mg/ml DAB and 0.024% H_2_O_2_ in PBS. Between each step, the membranes were washed three times for five minutes each with PBS/T buffer.

### 2.4. Analysis of gel- and membrane-based data using LC-MS/MS

Obtained gels and membranes were analyzed in Image Master 2D Platinum 7 Software (GE Healthcare, Chicago, IL, USA) to look for qualitative and/or quantitative differences in abundance/antigenicity of some venom proteins between the samples. Spots that were consistently different on subsequent gels/membranes in amount (expressed as %Vol parameter in the software) or unique to any samples were considered significant. Selected spots were excised from gels, and after destaining, reduced with DTT at a final concentration of 5 mM for 30 minutes at 60°C. Subsequently, alkylation was performed by adding iodoacetamide (IAA) at a final concentration of 15 mM and incubating for 30 minutes at room temperature in the dark. Following this, the samples were subjected to trypsin digestion (100 ng of the enzyme) for 18h at 37 °C. To deactivate trypsin, trifluoroacetic acid (TFA) in LC-MS grade water was added to the samples to reach a final concentration of 0.1% (v/v). The samples then were sonicated for 5 min, centrifuged at 14 000 × g for 5 minutes and mixed with 0.1% (v/v) TFA in LC-MS grade water in 3:5 ratio prior to LC-MS/MS analysis.

LC separation was carried out using the Dionex Ultimate 3000 Nano system (Thermo Fisher Scientific, Waltham, MA, USA) equipped with Acclaim PepMap RSLC nanoViper C18 column (75 μm × 25 cm, 2 μm particles) (Thermo Fisher Scientific, Waltham, MA, USA). The separation was performed using a 30-minute ACN gradient (ranging from 4% to 60%, in 0.1% formic acid). The chromatograph operated in online mode and was coupled with the Q ExactivePlusOrbitrap mass spectrometer (Thermo Scientific, Waltham, MA, USA). The analysis followed a data-dependent acquisition mode, with survey scans recorded at a resolution of 70,000 at m/z 200 in MS mode and 17,500 at m/z 200 in MS2 mode. Further fragmentation was performed on the 15 most prominent peaks from each MS spectrum. Spectra were acquired in the positive ion mode within the scanning range of 300–2000 m/z. The higher energy collisional dissociation (HCD) ion fragmentation method was employed, with normalized collision energies set to 25.

Peak lists obtained from MS/MS spectra were identified using 3 search engines: X!Tandem (ver. 2015.12.15.2), MS-GF+ (ver. 2021.01.08), and DirecTag. However, for some spectra, combinations of the following search engines were also used to facilitate the process of identification: OMSSA (ver. 2.1.9) and Comet (ver. 2020.01. rev. 2); DirecTag and Novor (v1.05.0573); Mascot server. Protein identification was conducted against a concatenated target/decoy UniProtKB Serpentes database (release 12/2022). The decoy sequences were created by reversing the target sequences in SearchGUI. The identification settings were as follows: Trypsin, Specific, with a maximum of 2 missed cleavages; 10.0 ppm as MS1 and 0.02 Da as MS2 tolerances; fixed modifications: Carbamidomethylation of C (+57.0214 Da), variable modifications: Oxidation of M (+15.9949 Da), Acetylation of protein N-term (+42.0105 Da). Peptides and proteins were inferred from the spectrum identification results using PeptideShaker (ver. 2.0.5) [22]. Peptide spectrum matches (PSMs), peptides, and proteins were validated at a 1% false discovery rate (FDR) estimated using the decoy hit distribution. Proteins identified from at least two peptides having at least two PSMs each were considered significant.

The mass spectrometry proteomics data have been deposited to the ProteomeXchange Consortium via the PRIDE [23] partner repository with the dataset identifiers: PXD043647 (10.6019/PXD043647) for *shotgun* LC-MS/MS experiment and PXD043722 (10.6019/PXD043722) for 2DE-LC-MS/MS approach.

### 2.5. Serial dilution affinity and avidity ELISA

MaxiSorp plates (Thermo Scientific, Waltham, MA, USA) were used for the assays. Venom samples, dissolved in PBS, were added to each well at a concentration of 100 ng. The plates were then incubated overnight at a temperature of 4°C. To prevent nonspecific binding, the wells were blocked with a 5% (w/v) solution of skim milk in PBS/T (PBS containing 0.1% Tween-20) for a duration of 3 hours. To ensure uniform antivenom concentration, all antivenom samples were initially adjusted to a concentration of 1 mg/mL (respective dilution factors: Antivipmyn Africa – 8.67x; EchiTAb+ICP – 181.88x; SAIMR Polyvalent – 206.33x). Serial five-fold dilutions of such samples, ranging from 1:100 to 1:1 562 500, were prepared. Each dilution was added to the respective wells, and the plates were incubated overnight at 4°C. Subsequently, a secondary anti-horse antibody conjugated with HRP was added to the wells at a dilution of 1:1000 in PBS/T. Substrate solution was prepared by combining 1 mg/mL TMB in DMSO with a sodium citrate-phosphate buffer (pH 5.0) at 1:9 ratio. Prior to signal detection, hydrogen peroxide was added to the substrate solution at a final concentration of 0.024%, and the mixture was added to the wells. The enzymatic reaction was allowed to proceed for 6 minutes, after which the absorbance was measured at 655 nm using Varioskan LUX microplate reader (Thermo Scientific, Waltham, MA, USA). Between each step, the wells were washed six times with PBS/T to remove any unbound substances. In addition to venom samples, control wells were included in the assay. These controls consisted of wells without venom as well as wells containing native horse antibodies without specific specificity, serving as the primary antibodies. The avidity ELISA test was conducted in a similar manner only for 1:500 dilution of primary antibody, with the exception that after the incubation step with primary antibodies, ammonium thiocyanate (at concentrations ranging from 0M to 8M) in water was employed to assess the binding strength of antibodies to antigens. The subsequent protocol remained unchanged. The statistical analysis was performed using GraphPad Prism 9 software. For the results obtained from the affinity ELISA, a two-way analysis of variance (ANOVA) with Tukey’s multiple comparison test was applied. p-values below 0.05 were considered statistically significant. For the data obtained from the avidity ELISA, individual data points were not compared directly. Instead, the slopes of the linear regression lines were compared.

## 3. Results

### 3.1. Proteomic analysis

#### 3.1.1. Shotgun LC-MS/MS analysis

The analysis of *Naja mossambica* venoms from different regions of Africa (Eswatini, Limpopo, and KwaZulu-Natal) using discovery *shotgun* LC-MS/MS provided detailed insights into the venom composition and revealed notable intraspecies differences. The data in Fig 1 shows general protein percentage profiles of venoms from distinct regions of Africa.

**Fig 1.**
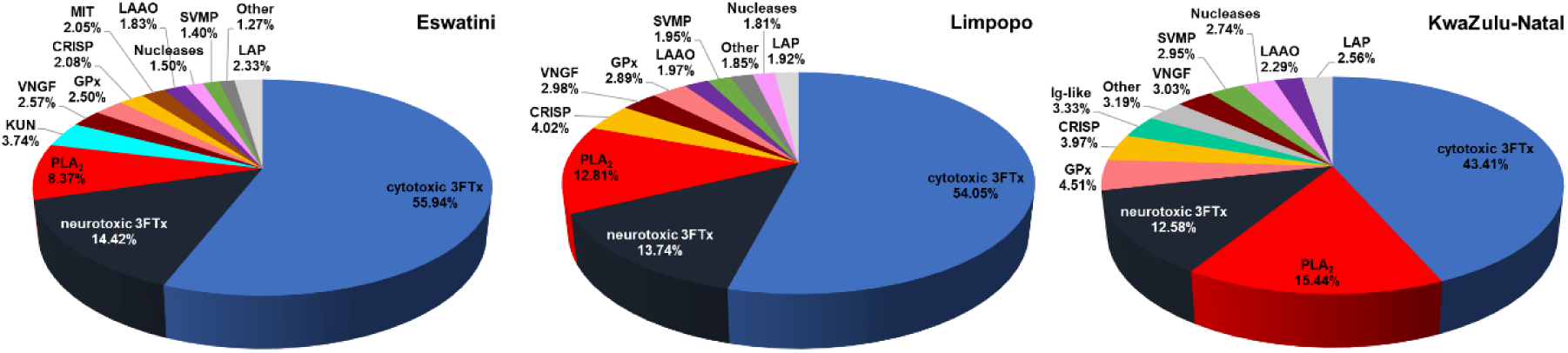
Pie charts showing the percentage of each protein family in *Naja mossambica* venoms from the Eswatini, Limpopo, and KwaZulu-Natal areas. The abbreviations used: 3FTx – three-finger toxin, PLA_2_ – phospholipase A_2_, KUN – Kunitz peptide, SVSP – snake venom serine protease, SVMP – snake venom metalloprotease, LAAO – L-amino acid oxidase, CRISP – cysteine-rich secretory protein, MIT – mamba intestinal toxin 1, VNGF – venom nerve growth factor, GPx – glutathione peroxidase, Ig-like – immunoglobulin-like protein, LAP – low abundant proteins. The list of all identified proteins is available in S1 Table.

In the KwaZulu-Natal group, the relative abundance of cytotoxic 3FTx was significantly lower compared to the other groups (around 12%). Conversely, both the KwaZulu-Natal and Limpopo groups exhibited approximately 4-7% higher amount of phospholipase A_2_ (PLA_2_) compared to the Eswatini group. Nevertheless, the most prominent protein groups in the venom of spitting cobras, 3FTxs and PLA_2_, collectively accounted for a major proportion of the tested venoms. The joint percentages of these two groups, however, differed substantially among the regions, with Eswatini venom comprising 78.73%, and Limpopo – 80.60%, while KwaZulu-Natal venom contained only 71.43% of these proteins. These discrepancies highlight substantial differences in venom composition and could impact potential variations in venom toxicity, immunogenicity, antigenicity, and biological activity.

Notably, a significant proportion of the identified three-finger toxins are proteins with neurotoxic activity. Moreover, a comparison of overall neurotoxin content, including neurotoxic 3FTx, Kunitz peptides, vespryns, and mamba intestinal toxin 1 (MIT) among the venom samples from different regions indicated that the venoms from Eswatini region exhibited a notably higher percentage (20.70%) of proteins with neurotoxic activity compared to the Limpopo (13.91%) and KwaZulu-Natal (12.80%) regions (Fig 1, S1 Table). Interestingly, Kunitz toxins accounted for nearly 4% of all venom proteins in the samples from Eswatini which may suggest their non-negligible influence on the venom biological effect.

A notable amount of glutathione peroxidase protein (GPx) was identified in all groups, comprising a significant percentage (2.5 – 4.5%) of the total proteins in the tested venoms. It is an unexpected result as GPx has been classified as a rare protein family and is usually identified in a minor amount [24]. Within the KwaZulu-Natal group, a considerable proportion of proteins from the Immunoglobulin-like (Ig-like) group were detected (∼3.5%), consistent with our previous reports for different venoms from *Naja* genus, where this group of proteins was also identified [15, 25–26]. There is also a significant difference in SVMP content between samples indicating the highest amount of these enzymes in venoms from KwaZulu-Natal region.

To further investigate the intraspecies differences in *Naja mossambica* venoms, statistical analysis of LC-MS/MS data between tested samples was performed, providing additional insights into the venom composition of the studied African regions.

PCA analysis of the quantitative MS data for the identified proteins in the three different venom samples revealed distinct grouping patterns (Fig 2a). The samples originated from the same region formed homogeneous clusters, indicating the presence of specific venom profiles within the same species. Clear differences were observed between the groups, further highlighting the unique characteristics of the venoms originating from different regions.

**Fig 2.**
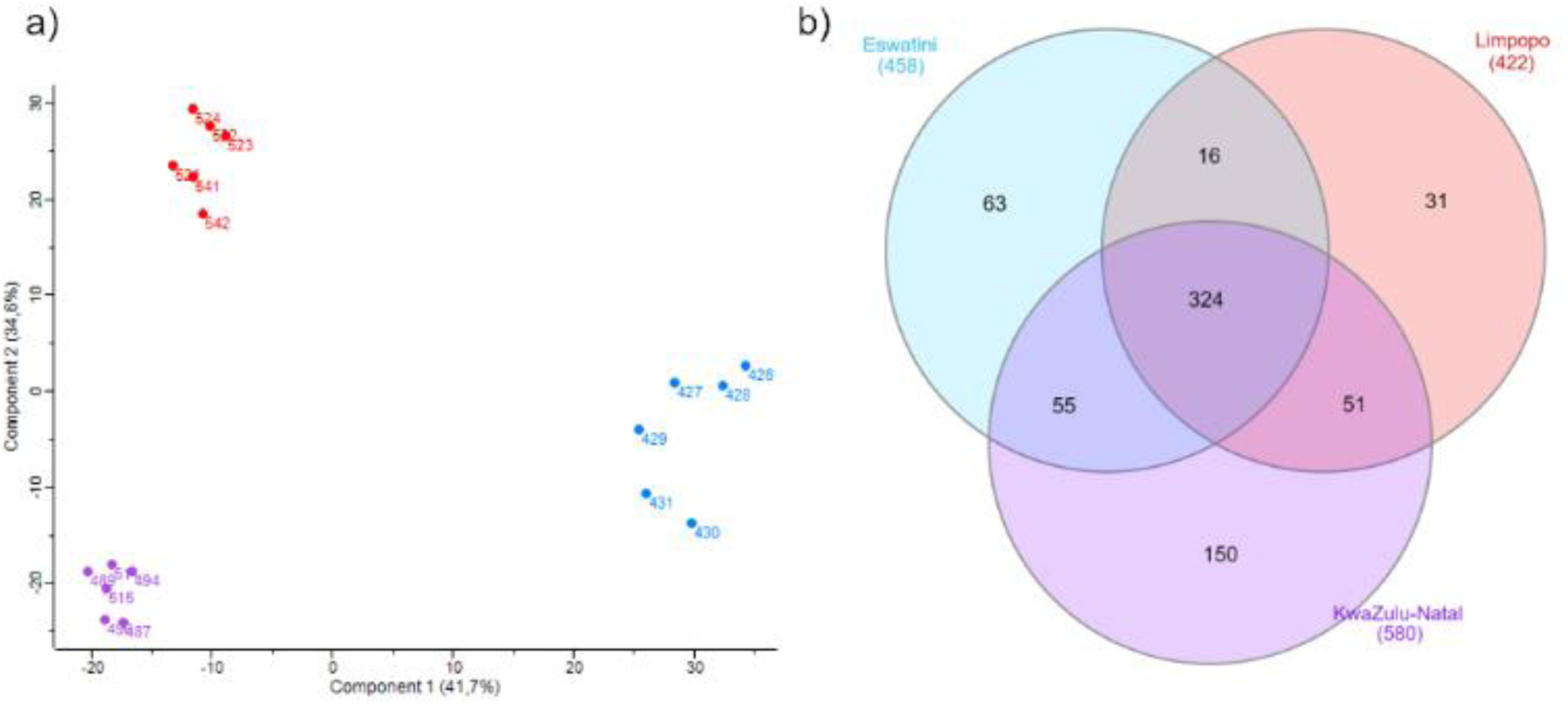
**a)** Principal Component Analysis (PCA) of the quantitative MS data for proteins identified in venoms of different *Naja mossambica* populations. The points on the graph indicate individual venom samples (blue - Eswatini; red - Limpopo; purple - KwaZulu-Natal); **b)** Venn diagram showing the number of shared and unique peptides identified in *N. mossambica* venom samples from different regions of Africa. Data for peptides that were identified in at least three samples from certain group. All identified peptides are listed in S3 Table.

Moreover, the analysis of shared and unique peptides provided valuable insights into the composition of the venom samples (Fig 2b). The venom samples from the KwaZulu-Natal region exhibited the highest number of identified peptides. A total of 150 unique peptides were exclusively detected in samples from this particular region, which were not found in the venom samples from the other two populations.

To further investigate the LC-MS/MS data, hierarchical clustering was performed and heatmap was generated showing the proteins whose presence/quantity significantly differs between venoms collected from snakes belonging to populations that inhabit distinct areas of Africa (Fig 3).

**Fig 3.**
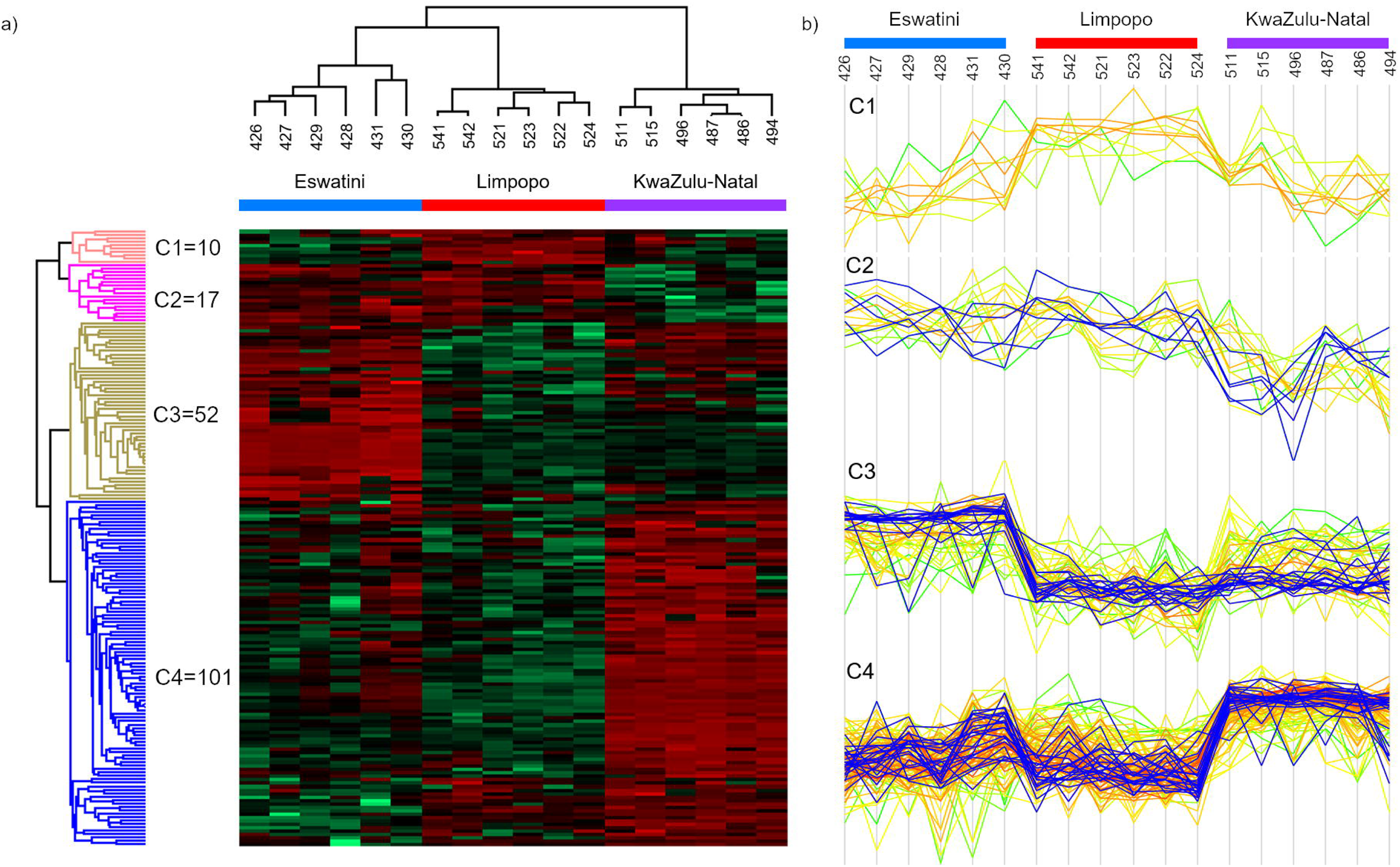
Data show relative abundance of proteins that differs in quantity between venom samples from distinct parts of Africa. A total of 180 proteins were subjected to hierarchical clustering analysis and **a)** visualized in a heatmap. The column tree reveals distinct protein profiles for samples obtained from various regions. The heatmap utilizes red and green colors to indicate higher and lower protein abundance, respectively. Based on the clustering analysis, four clusters were identified, representing proteins with similar quantity profiles across the samples. **b)** Profile plots corresponding to each cluster are displayed on the right side of the figure. Enriched proteins within each cluster are highlighted in blue. For a comprehensive list of these proteins, refer to the S4 Table.

The resulting heatmap revealed four distinct clusters, characterized by variations in the content of specific proteins across the samples. Cluster 1 is the only group of proteins whose amount is increased in Limpopo venom. This cluster consisted of 10 proteins without a clear family trend. Notably, among these proteins, there was one neurotoxic 3FTx (Short neurotoxin 3; P01432) that exhibited higher abundance in all tested Limpopo samples (S3 Table).

There are 17 proteins in the second cluster (C2), the amount of which is higher in the samples from Eswatini and Limpopo compared to the samples from KwaZulu-Natal. In this case, the annotation enrichment analysis showed that proteins belonging to the ‘Snake cytotoxin family, cobra-type’ (IPR003572), are overrepresented in this group (S3 Table). The quantitative trend of these proteins is marked in blue colour in Fig 4b (Cluster C2). Of the five proteins classified within this family, four are cytotoxins which aligned with the overall lower content of cytotoxic three-finger toxins observed in the KwaZulu-Natal group (Fig 1).

**Fig 4.**
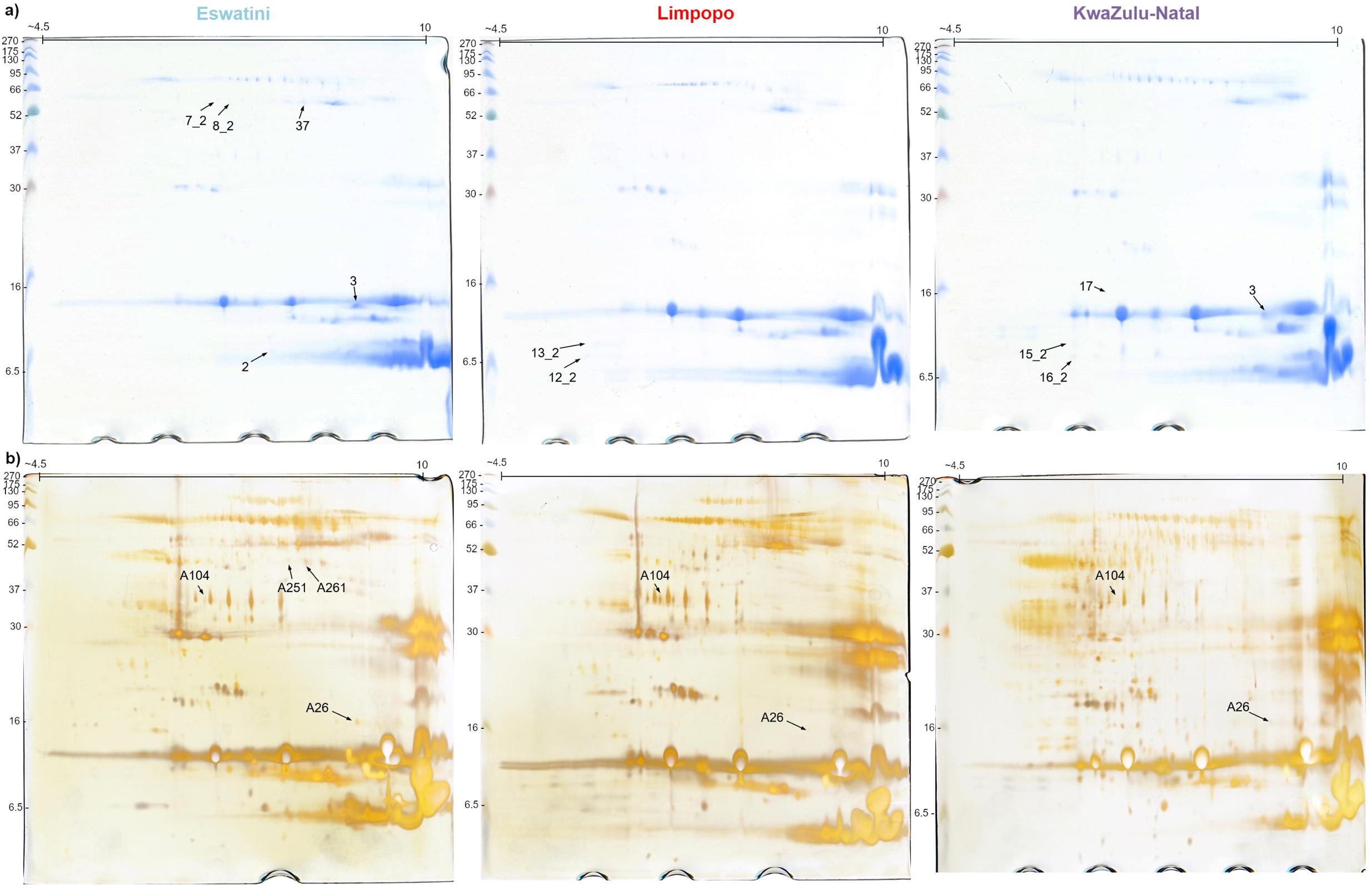
Two-dimensional protein maps of venom samples from distinct regions of Africa (Eswatini, Limpopo, KwaZulu-Natal). **a)** gels stained by Coomassie Brilliant Blues G-250 **b)** gels stained with silver. Marked spots represent successfully identified proteins that are listed in Table 1.

The next cluster (C3) comprised proteins that were significantly more abundant in the Eswatini venom comparing to other venoms. Within this cluster of 52 proteins, 16 corresponded to keywords such as ‘Ion channel impairing toxin’, ‘Neurotoxin’, and ‘Postsynaptic neurotoxin’ in the UniProt database. Specifically, 11 proteins are classified as neurotoxic 3FTxs, three as Kunitz peptides, and two as vespryns. Additionally, among the remaining 36 proteins, four other neurotoxins (including two 3FTxs, one KUN, and one MIT) were found, but since they are not sufficiently annotated in the UniProt database, they did not group during the enrichment analysis (S3 Table).

Cluster 4, the largest cluster, encompassed 101 proteins that exhibited significantly higher abundance in the venom samples from the KwaZulu-Natal region. Annotation enrichment analysis identified a substantial overrepresentation of proteins belonging to the ‘Immunoglobulin-like domain superfamily’ (IPR 036179). Out of the 24 proteins assigned to this superfamily, 21 were assigned to the fourth cluster, highlighting the distinct nature of the KwaZulu-Natal samples in terms of this protein group (S3 Table).

In addition to Ig-like proteins, other protein groups with unknown function in venom can be found within this cluster, including Ezrin-like and Cobalamin-binding intrinsic factor (CBLIF) groups. The data on the heatmap indicate an increased presence of these proteins in venoms from the KwaZulu-Natal region. For a comprehensive list of proteins categorized into individual clusters, refer to the S3 Table.

#### 3.1.2. Two-dimensional electrophoresis

Intraspecies differences in *N. mossambica* were also analyzed using two-dimensional electrophoresis (2DE) with two different gel staining protocols, CBB G250 (Fig 4a) and silver staining (Fig 4b).

Silver staining allowed for the visualization of a larger number of proteins compared to CBB staining clearly presenting exceptional complexity of snake venoms. This enhanced sensitivity was particularly advantageous in capturing subtle differences in venom profiles, however, obtaining high reproducibility in the 2DE approach was difficult and thus, in Fig 1, only spots that could be unequivocally selected as differing and successfully identified by MS are labeled. Nevertheless, the results allowed to reveal some variations in venom composition among the African regions. The identified spots are labeled with corresponding numbers on the gels, and detailed information regarding their identification can be found in Table 1.

**Table 1.**
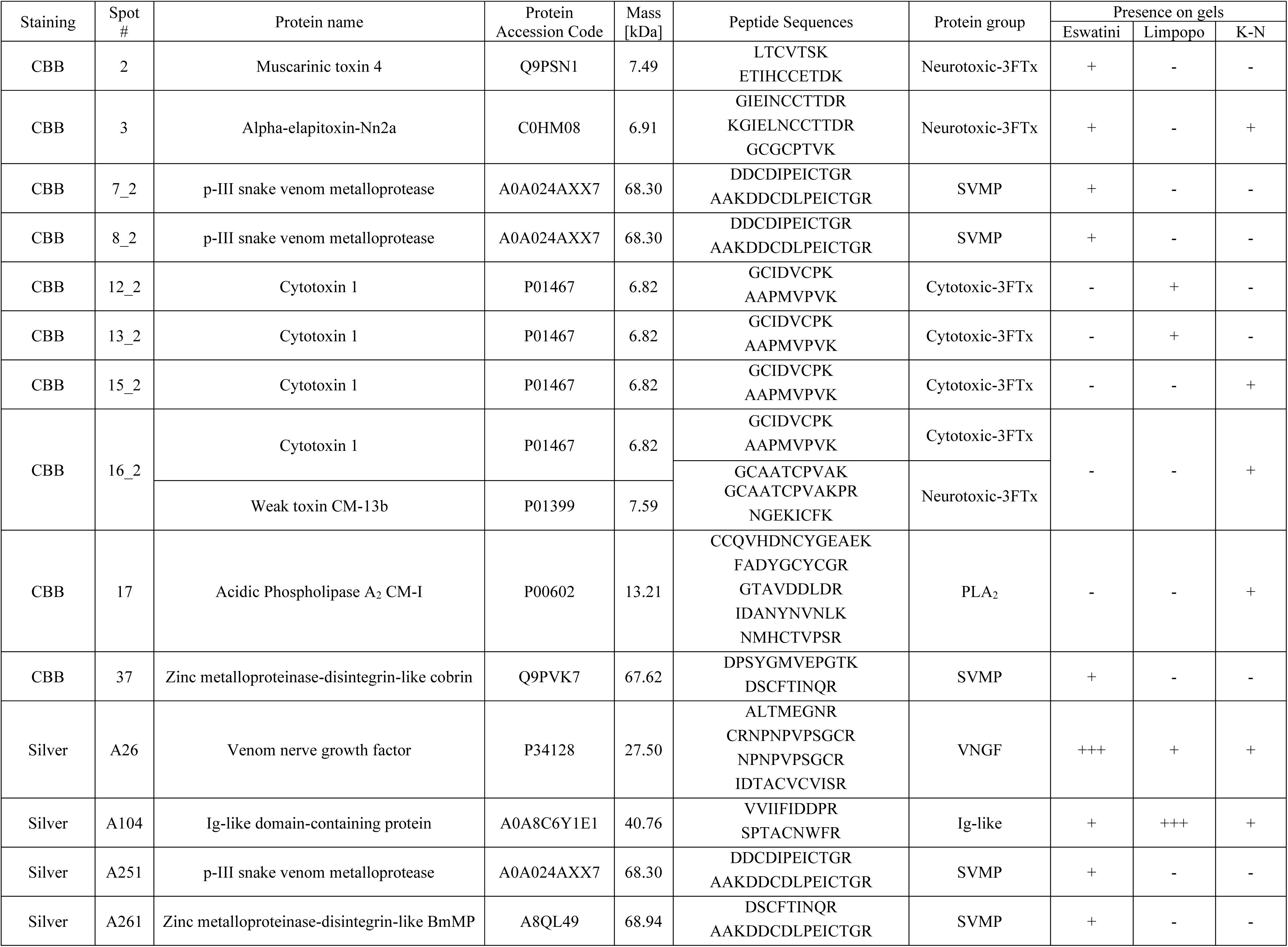
The list of identified proteins by 2DE-LC-MS/MS approach (+++ higher abundance on gel + presence on gel; - absence on gel)

In general, the 2D gels obtained for samples from the KwaZulu-Natal region exhibited the highest number of spots, indicating a more diverse range of venom components which is consistent with LC-MS/MS data presented on the Venn diagram (the highest number of identified peptides) (Fig 2b). Moreover, proteins clustered in the 3FTx region appeared to be less numerous in the KwaZulu-Natal venom gels compared to the other regions. This observation is also supported by the data obtained by the direct shotgun MS approach (Figs 1 and 3). This clearly indicates potential variations in the abundance of these proteins among tested populations of *Naja mossambica*.

A large and visible spot (#3) was found to be present in the Eswatini and KwaZulu-Natal gels but was absent in the Limpopo gels. Through further analysis, this protein was identified as a neurotoxin belonging to the short-chain subfamily of 3FTx. Limpopo gels exhibited also additional signals in the middle area, which were identified as Ig-like proteins (#104) and are probably ancillary proteoforms of these proteins. On the other hand, Eswatini gels displayed unique spots in the ∼45 kDa area, two of which were identified as proteins from the SVMP family (#A251, #A261).

Despite the predominant clustering of proteins from the 3FTx group in one area, several different spots across the gels contained proteins belonging to this family. However, due to limitations in the bottom-up method and incomplete databases, obtaining precise information about the protein sequences and specific proteoforms was challenging. Thus, it was just established that these proteins were part of the 3FTx family, as indicated by spot labels (#12_2, #13_2, #15_2, #16_2).

### 3.2. Immunochemical analysis

#### 3.2.1. Two-dimensional Western Blot

Proteomic analyses allowed us to show general differences in the composition of the venoms collected from different *N. mossambica* populations. To explore the differences in venom-antivenom interactions between the samples we used a two-dimensional Western Blot approach combined with LC-MS/MS for selected spots (Figs 5 and 6; Table 2). This enabled us to investigate and compare the differential binding patterns between the venoms and the three tested antivenoms (EchiTAb+ICP, Antivipmyn Africa, and SAIMR polyvalent). We utilized membranes with distinct pore sizes (0.2 and 0.45 μm) to allow the binding of both high- and low-molecular-weight proteins.

**Fig 5.**
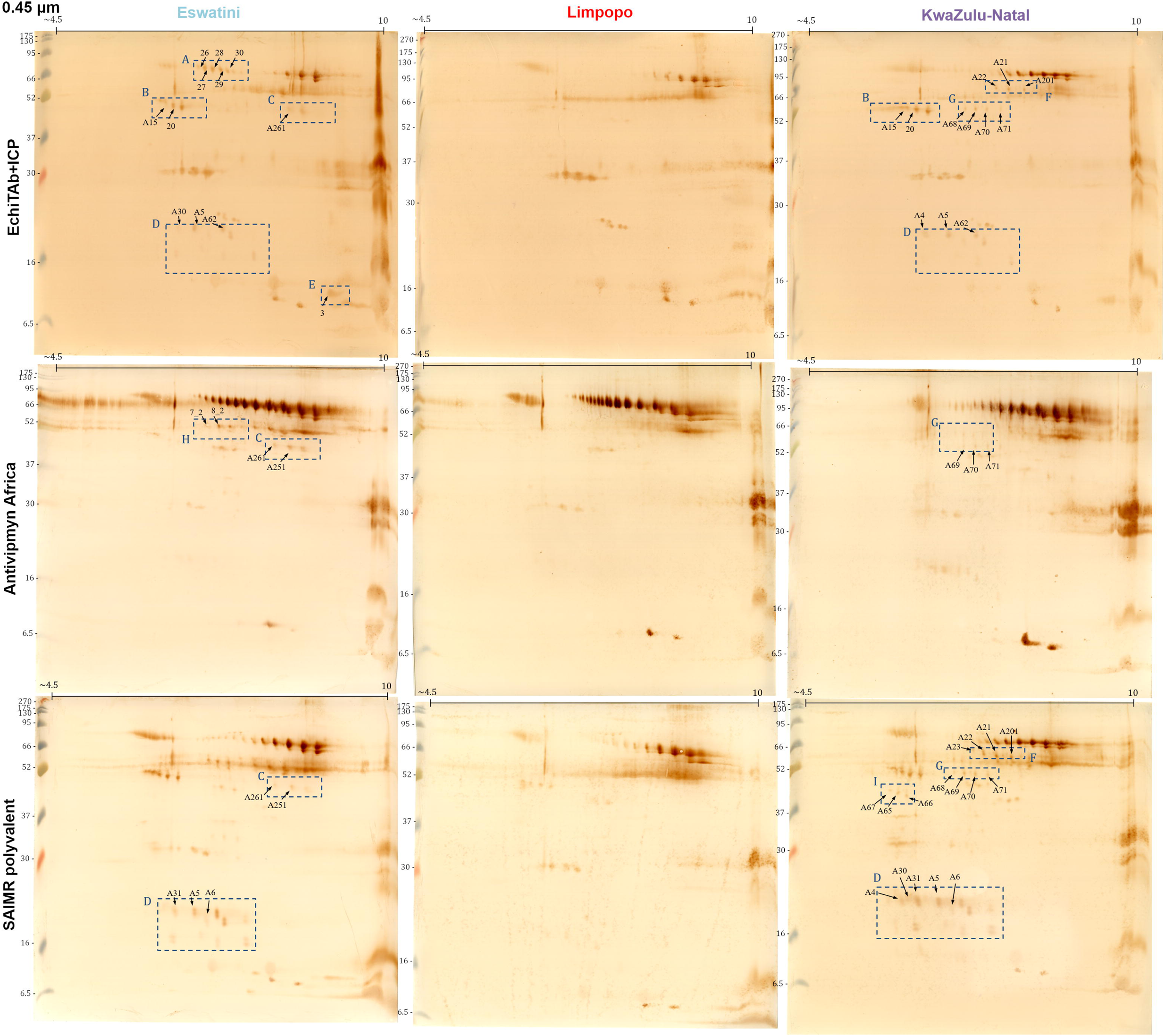
Two-dimensional Western Blots of three venom samples (Eswatini, Limpopo, KwaZulu-Natal) with the use of three different antivenoms (EchiTab+ICP, Antivipmyn Africa, SAIMR Polyvalent) as primary antibodies performed on 0.45 μm nitrocellulose membranes. Dashed, blue squares indicate areas that differ in signal pattern between venoms. Marked spots represent successfully identified proteins that are listed in Table 2.

**Fig 6.**
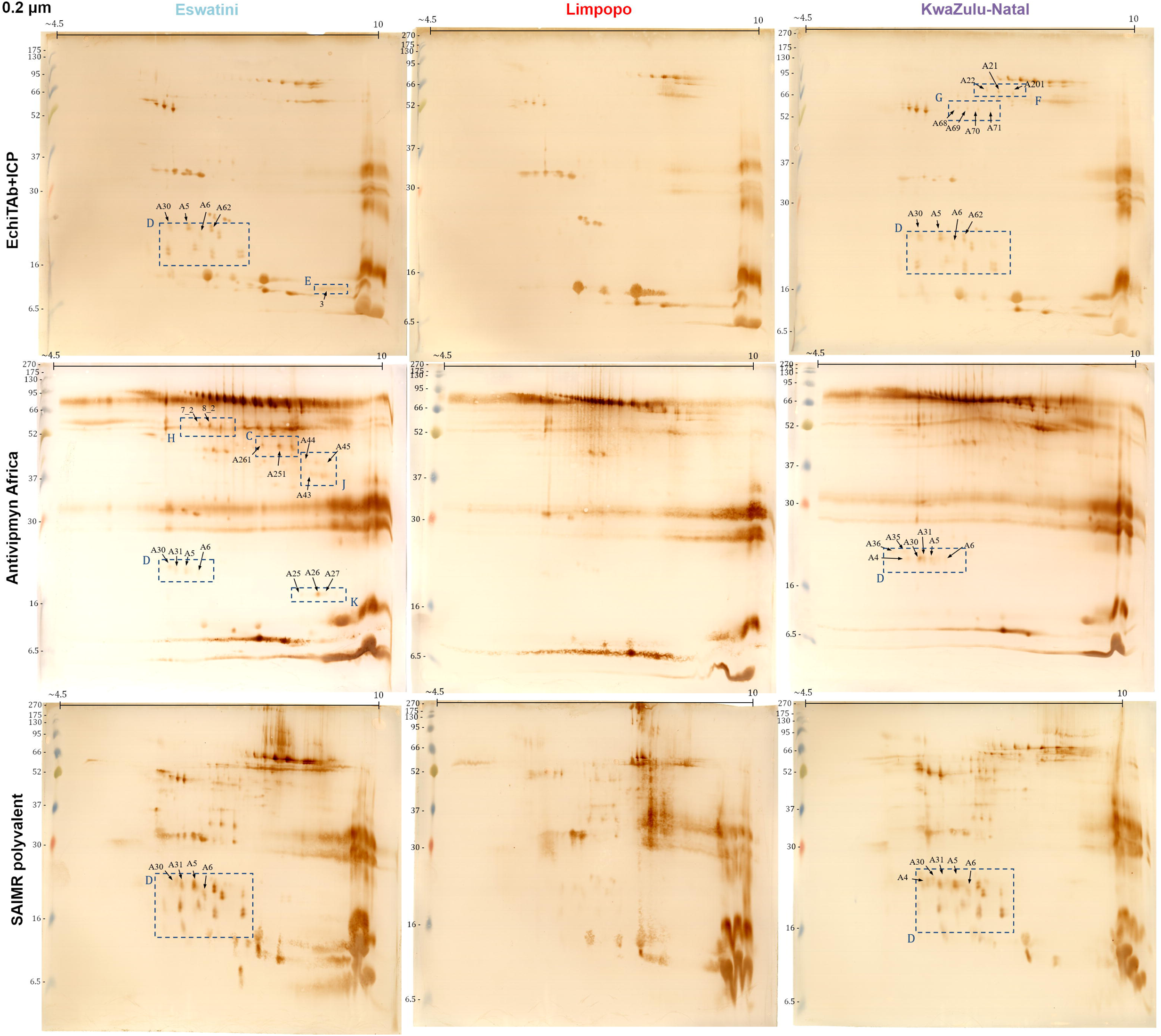
Two-dimensional Western Blots of three venom samples (Eswatini, Limpopo, KwaZulu-Natal) with the use of three different antivenoms (EchiTab+ICP, Antivipmyn Africa, SAIMR Polyvalent) as primary antibodies performed on 0.2 μm nitrocellulose membranes. Dashed, blue squares indicate areas that differ in signal pattern between venoms. Marked spots represent successfully identified proteins that are listed in Table 2.

**Table 2.**
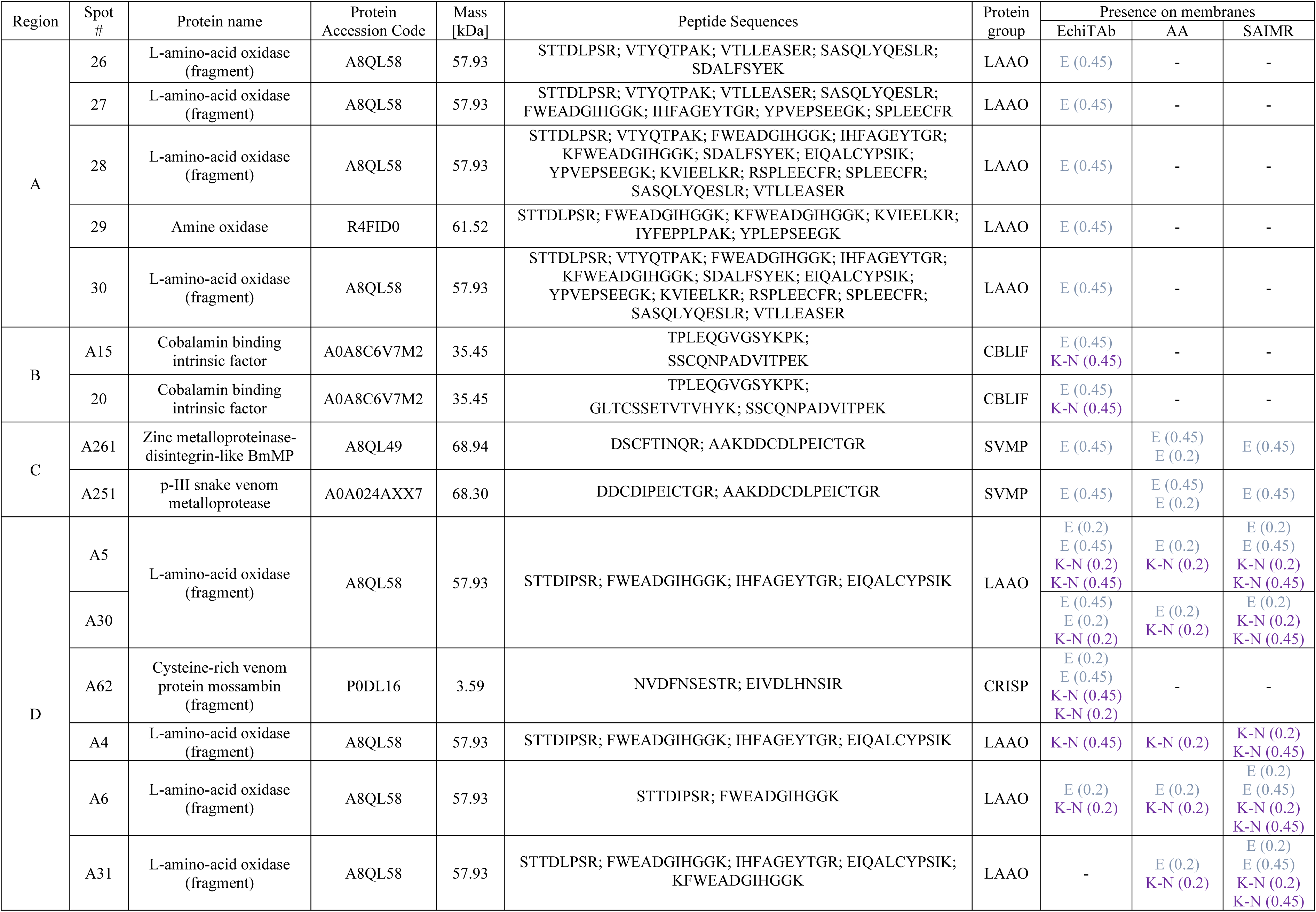
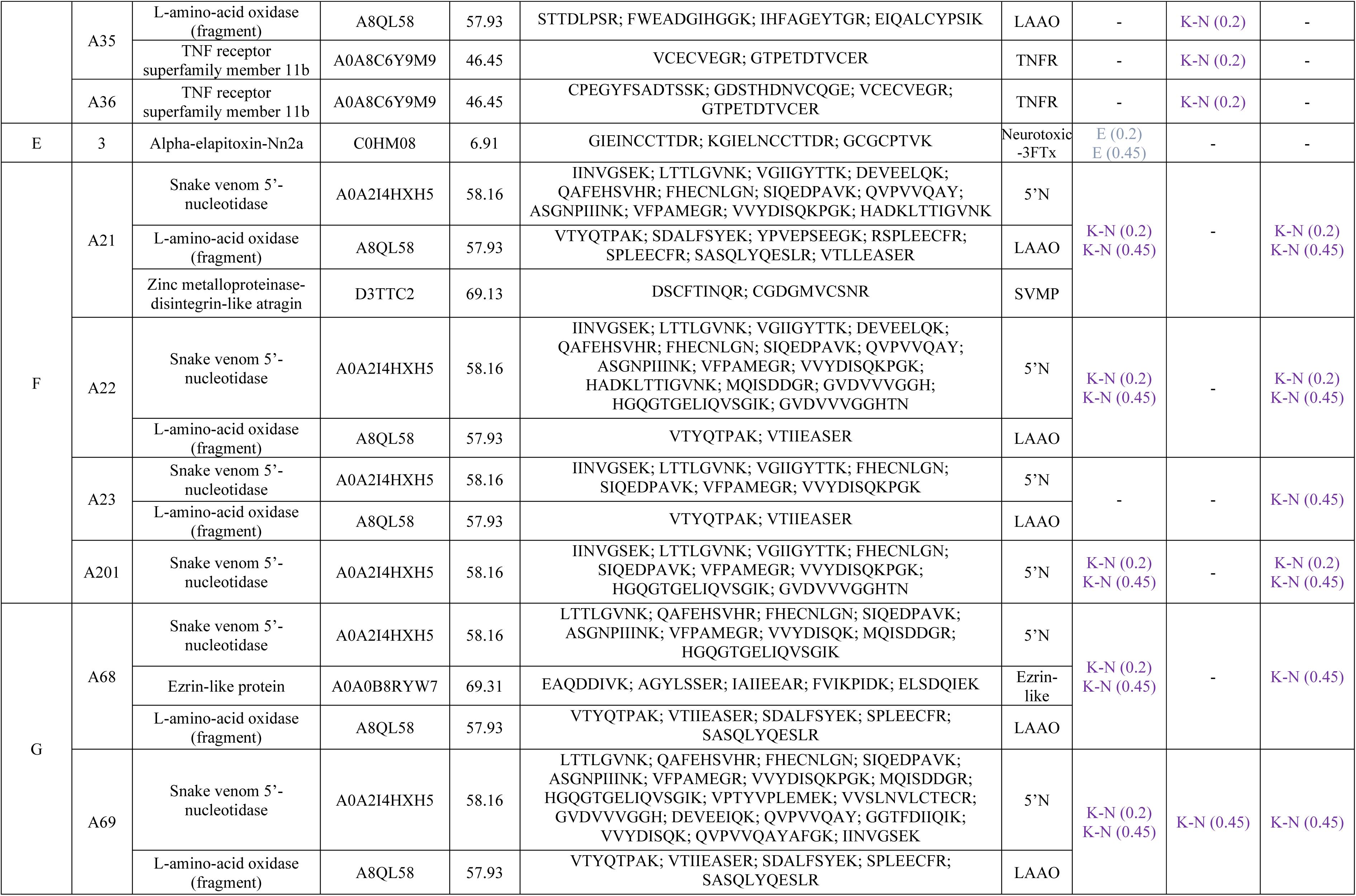
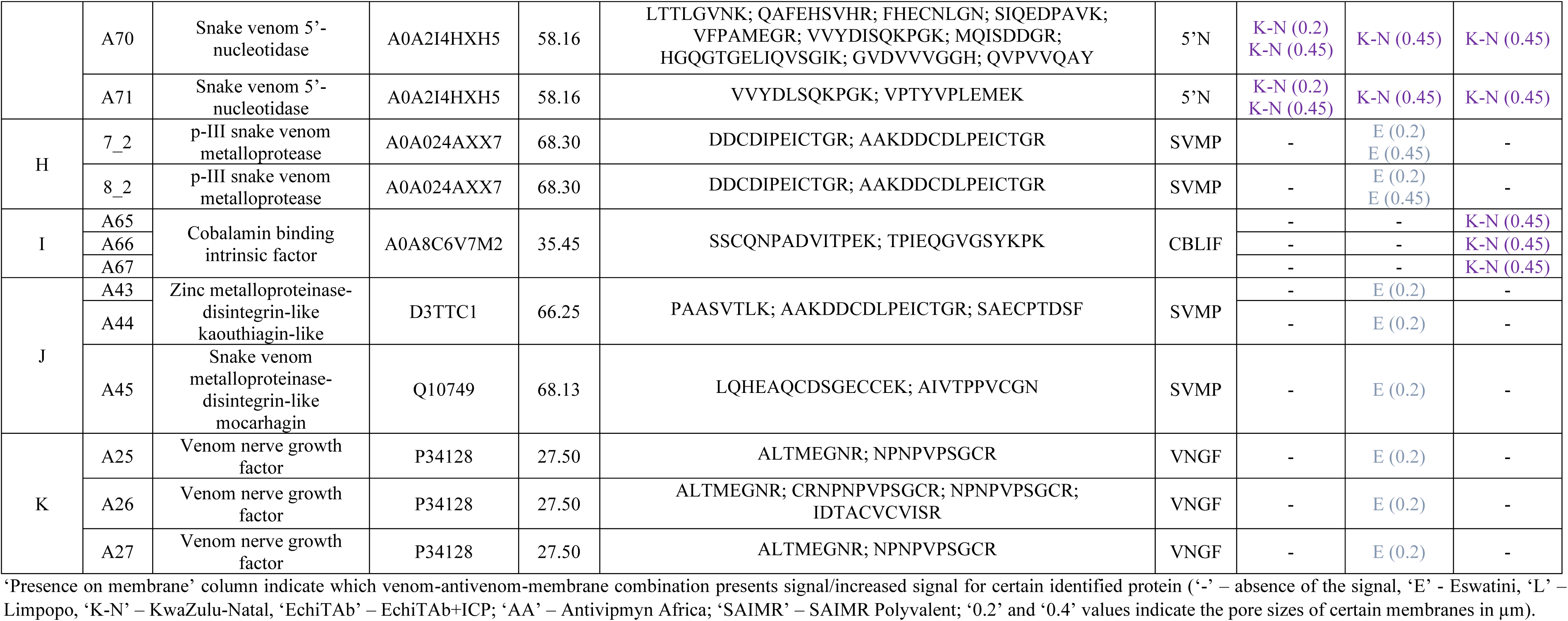
The list of proteins selected based on 2D Western Blot analysis and identified by 2DE-LC-MS/MS approach.

Membrane analysis highlighted 11 areas (A-K) showing a unique pattern of signals that differed between the tested venoms in a qualitative and/or quantitative manner. The proteins marked with numbers on the membranes were successfully identified using mass spectrometry, enabling their characterization.

At first, 2D Western Blot analyses clearly demonstrated the lowest signal complexity for samples originating from the Limpopo region. Moreover, significant differences in protein interactions were observed in areas A-G of the membranes with the use of EchiTAb+ICP antivenom. Areas A, C, and E exhibited characteristic signals associated with LAAO, SVMP, and neurotoxic 3FTx proteins, respectively, which were more prominent in venom from the Eswatini region. Conversely, regions F, G, and I which contained several distinct protein groups, were characteristic of venom from the KwaZulu-Natal region. The presence of Ezrin-like and CBLIF proteins was exclusive for the KwaZulu-Natal membranes what is consistent with proteomics MS data observed in cluster C4 in Fig 3.

Moreover, proteins identified as Cobalamin-binding intrinsic factor (B, I) differentiated the membranes obtained using EchiTAb+ICP and SAIMR antivenoms, indicating variations in the interaction patterns of venom proteins with these different antivenoms. On the other hand, Antivipmyn Africa antivenom was especially effective in distinguishing areas H, J (both SVMPs), and K (VNGF), which were characteristic of venom from the Eswatini region.

Among the different areas, the D region stood out as the most distinctive in terms of the interaction of venom proteins with all tested antivenoms. Signals within this area were consistently absent or very weak for the venom samples from Limpopo. Interestingly, eight proteins from three different protein families (LAAO, CRISP, TNFR) were successfully identified within this area. Five spots represent proteins from the LAAO family, which were also detected in the area A. Moreover, two spots within this region contain peptides assigned to TNF receptor family proteins of unknown function in a venom. These findings from the two-dimensional Western Blot analysis provide interesting insights into the differences in protein reactivity and expression profiles among *Naja mossambica* venoms from the studied regions, highlighting distinct patterns of venom composition and antigenicity.

In addition, the analysis of the Western Blot membranes for different antivenoms revealed interesting differences in their overall appearance and reactivity with venom proteins. These observations shed light on the potential variations in antigen recognition and binding capacities between tested antivenoms. Antivipmyn Africa antivenom exhibited a distinct pattern on the membranes, suggesting a higher potential for interaction with high-molecular-weight venom proteins compared to the other antivenoms. However, it was observed that this antivenom yielded weak or no signals in the D area, which emerged as a crucial differentiating factor between samples from Eswatini and KwaZulu-Natal regions compared to Limpopo. Furthermore, the utilization of EchiTAb+ICP and SAIMR antivenoms resulted in the appearance of signals in similar areas on the membranes. However, the overall signal intensity was notably higher on the membranes obtained using SAIMR polyvalent antivenom.

#### 3.2.2. Quantitative analysis of the venom-antivenom interaction with affinity and avidity ELISA

Quantitative assessment of the interaction between venom toxins and tested antivenoms was also conducted using serial dilution affinity and avidity ELISA assays (Figs 7 and 8).

**Fig 7.**
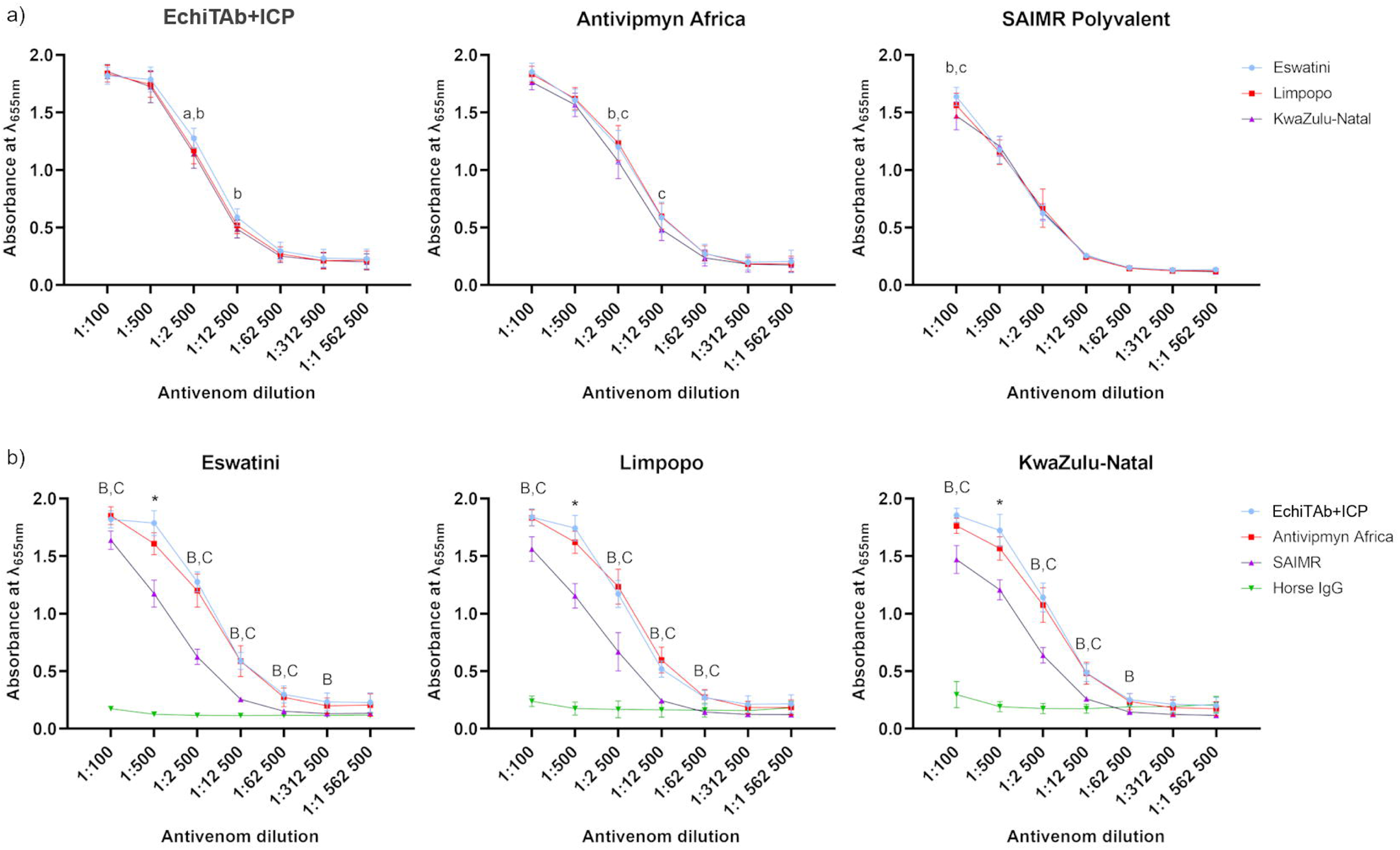
Serial dilution affinity ELISA results show quantitative data on venom-antivenom interactions represented as **a)** each antivenom against all three tested venoms and **b)** each venom against all tested antivenoms. Data points are the mean of absorbance values at 655 nm from three technical replicates with error bars representing standard deviations. The significance of mean differences was tested by two-way ANOVA with Tukey’s test for multiple comparisons (α = 0.05). Letters or symbols above the error bars indicate which groups are different from each other (a – Eswatini and Limpopo; b – Eswatini and KwaZulu-Natal; c – Limpopo and KwaZulu-Natal; A – EchiTAb+ICP and Antivipmyn Africa; B – EchiTAb+ICP and SAIMR polyvalent; C – Antivipmyn Africa and SAIMR polyvalent; * - all samples are different)

**Fig 8.**
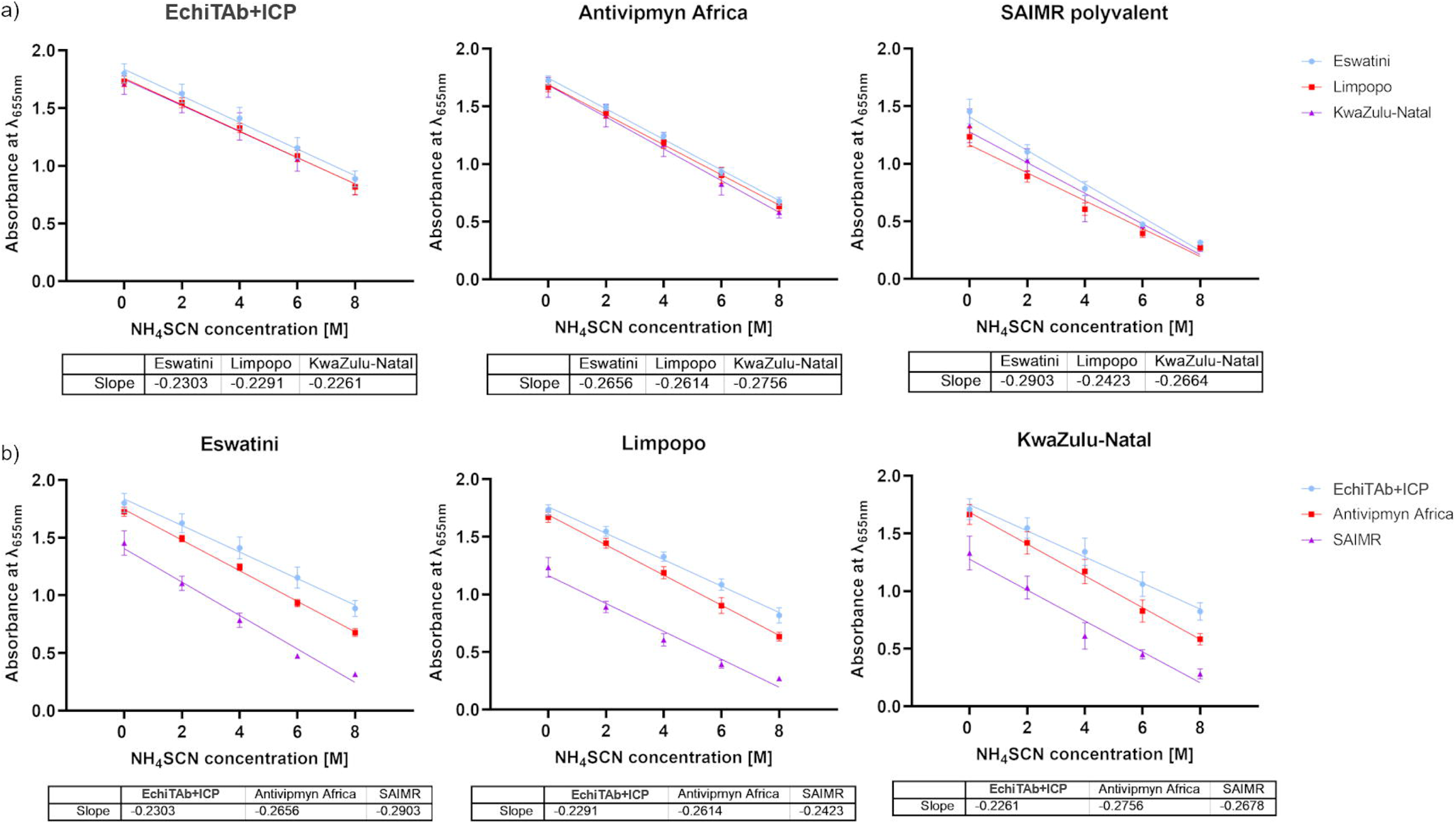
Avidity ELISA results show venom-antivenom interaction strength in the presence of ammonium thiocyanate chaotrope. The primary antibodies (antivenoms) in the experiment were used in 1:500 dilution. The data were graphed to show **a)** the difference in avidity between different venoms for each tested antivenom; **b)** the difference in avidity between each antivenom for any given venom tested. Data points are the mean of absorbance values at 655 nm from three technical replicates with error bars representing standard deviations. Least squares linear regression was used to fit the lines into the experimental data points. The tables below each graph contain numeric values for each slope of the lines.

The objective of the serial dilution affinity ELISA experiment was twofold: to quantify the reactivity of the tested venoms with each antivenom (Fig 7a) and to compare the general efficiency of venom protein binding by the various antivenoms (Fig 7b). The results revealed subtle differences in the overall interactions between venoms from the three distinct African regions and the respective antivenoms (Fig 7a). Obtained data indicate that at a dilution of 1:2500 for the EchiTAb+ICP, venom from Eswatini exhibited the highest reactivity, while for the Antivipmyn Africa antivenom, the lowest reactivity has been shown for venom from the KwaZulu-Natal region. The SAIMR polyvalent showed distinguishable differences in reactivity only at the 1:100 dilution, with the venom from the KwaZulu-Natal region exhibiting the lowest signal.

On the other hand, significant disparities in the effectiveness of the individual antivenoms were observed (Fig 7b). Such data representation clearly shows that at a dilution of 1:500, significant variations in optical density (OD) were observed for all venoms across the tested antivenoms. Interestingly, the EchiTAb+ICP consistently exhibited the highest binding capacity for venom proteins, while the SAIMR polyvalent antivenom displayed the lowest reactivity against venom proteins. In this context, however, it is crucial to note that antivenom concentrations before the test were standardized to 1 mg/ml. Therefore, the graph can only be interpreted in the context of comparing the efficacy of toxin binding by the equal amount (same mass) of antibody molecules in each antivenom. SAIMR (dilution factor: 206.33x) and EchiTAb+ICP antivenoms (dilution factor: 181.88x) were significantly more diluted than Antivipmyn Africa (dilution factor: 8.67x) before the experiments. Hence, the effectiveness of these two former products would likely be much higher than Antivipmyn Africa in undiluted antivenoms due to a greater number of neutralizing molecules.

The results from the avidity ELISA assay did not reveal much new significant data in addition to previous results from the experiment estimating quantitatively venom-antivenom affinity (Fig 8).

The analysis of the avidity of the antibodies contained in the individual antivenoms confirmed the findings obtained from the affinity ELISA, reaffirming rather than introducing novel insights that would alter the interpretation of the prior results (Fig 8). There are no substantial differences in the avidity when comparing the data obtained for different venoms with the use of the same antivenom. The highest discrepancy in the slope of the line occurs between Eswatini (−0.2903) and Limpopo (−0.2423) with SAIMR antivenom which is not a big difference considering non-negligible values of SD (Fig 8a). The analysis of the slope values depicted in Fig 8b reveals subtle variations among the different antivenoms. The difference between antibody binding strength of EchiTab+ICP and SAIMR towards Eswatini venom proteins seems the most noticeable (slope values of −0.2303 and −0.2903 respectively), which could indicate a slightly lower avidity of SAIMR antivenom, however, the observed change is not statistically significant. On the other hand, once again, the OD values demonstrated the highest reactivity for the EchiTAb+ICP antivenom, while the SAIMR polyvalent exhibited significantly lower OD values (Fig 8b).

## 4. Discussion

In recent years, there has been an intensive search for alternatives to current therapeutic solutions following snakebites, including innovative approaches such as using recombinant antivenoms based on monoclonal antibodies, employing small molecules to inhibit the activity of certain enzymes, and utilizing virus-derived protein scaffold to neutralize venom toxins [27–29]. However, until new solutions are developed, snakebite treatments predominantly rely on the use of antivenoms obtained through the immunization of large domestic animals. Currently, in Africa, there are still approximately half a million snakebites annually, posing a significant challenge, especially considering the inadequate supply of effective antivenoms [3]. Therefore, beyond exploring new solutions, it is equally crucial to assess the existing therapies and enhance their efficacy [30].

In this paper we aimed to analyze the intraspecific differences in *Naja mossambica* venoms collected from three different regions in south Africa and their impact on the interactions with three commercial antivenoms. To achieve this, we have used a combination of proteomics and immunochemical approach.

Proteomics analysis brought interesting new data on complexity and variety of *Naja mossambica* venoms. In general, previously published proteomic data for *N. mossambica* venom are relatively consistent with those obtained in this study. The overall trend characteristic of *N. mossambica* venom, as well as other spitting cobras where the majority of venom comprises 3FTx proteins and phospholipases A_2_, was confirmed [14,16]. However, when it comes to the exact values obtained for these protein groups and other toxins, differences between the data are more apparent. Due to the limitations of quantitative mass spectrometry, especially for complex mixtures, and the use of different methodologies [26], our intention is not to compare the exact values to available literature data. Instead, we primarily focus on comparing data between the studied populations of *N. mossambica*, which were obtained using a uniform methodology applied in this study. The only quantitative values we would like to confront with previous literature reports are that concerning the neurotoxic 3FTx toxins found in *N. mossambica* venom. These findings present an obvious contrast to earlier data and could considerably impact the perception of this spitting cobra’s venom proteome. It is noteworthy that neurotoxic 3FTxs constitute almost 15% of all proteins in each of the analyzed venoms. This percentage is significantly higher than previously reported values of 1.6% [14] and 4.39% [16] for the proportion of these toxins in *N. mossambica* venom. Moreover, apart from *Naja nubiae* venom, where the percentage of neurotoxins slightly exceeded 10% of all venom proteins, no other spitting cobra venoms registered neurotoxic 3FTxs in amounts greater than 5% [14, 16]. Furthermore, considering other proteins that can act on the nervous system, such as Kunitz toxins, vespryns, and the MIT1 protein, which together constituted over 6% of all venom proteins from Eswatini region, these findings suggest that the venoms of these snakes could exert various neurotoxic effects. A more detailed quantitative analysis of identified neurotoxic 3FTxs in analyzed venoms shows that 72 to 86% of them were short-chain 3FTxs, 11-23% were weak toxins, and 3-5% were long-chain 3FTxs (S3 Table). Among the identified proteins were α-neurotoxins, calciseptins, fasciculins, mambalgins, and toxins interacting with adrenergic and/or muscarinic receptors (S3 Table). Therefore, the significantly higher neurotoxin content, and their diversity observed in the Eswatini venoms (notably, the exclusive identification of the MIT1 neurotoxic protein from the prokineticin family and higher number of identified proteins from KUN and 3FTxs families in this venom), may play a pivotal role in shaping a distinct clinical profile for individuals envenomed by snakes from this region and could influence their treatment outcomes. This trend is further depicted in Fig 3, where proteins with neurotoxic activity clearly grouped within cluster 3, which contains proteins more abundant in Eswatini venoms than in the other samples.

Another cluster that effectively differentiates analyzed venom samples is cluster C4. This cluster is formed by the proteins markedly increased in KwaZulu-Natal venom samples, and it includes 21 proteins belonging to the ‘Immunoglobulin-like domain superfamily’ (IPR 036179). While these proteins are not officially recognized as known venom components [24], our previous reports have detected them [15, 25–26]. This analysis indicates they constitute nearly 4% of all venom proteins in the KwaZulu-Natal venom and around 0.6-0.7% in two others. There are also other reports documenting the identification of Ig-like domain-containing proteins [31–32]. However, their presence in the venom was not discussed, likely because they do not belong to recognized toxin groups/families. Nevertheless, given the substantial number of identified peptides mapping to snake proteins within a single InterPro database entry (IPR036179), we believe they represent uncharacterized proteins that contain the Ig-like domain. It is worth noting that C-type lectins and zinc metallopeptidases, recognized toxins from snakes’ venom, may possess similar domains in their structure in other organisms [33–35].

To compare venom proteomes from different regions, a two-dimensional electrophoresis technique with protein identification by LC-MS/MS was also applied. In this case, it is noteworthy that gels obtained for KwaZulu-Natal venoms showed the highest complexity, especially visible on silver-stained gels (Fig 4b), which corresponds to the data obtained during shotgun LC-MS/MS analysis (Fig 3). On the other hand, KwaZulu-Natal gels revealed lower signal intensity than other venoms in the bottom right region, corresponding to the presence of small and basic toxins like 3FTx proteins, which also aligns with shotgun LC-MS/MS data (Fig 1). Additionally, in relation to proteins containing the Ig-like domain, a member of this group was identified in spot #A104, with a monomer mass of approximately 37 kDa.

We also used affinity and avidity ELISAs to measure and compare how reactive various *N. mossambica* venoms were with three different antivenoms: EchiTAb+ICP, Antivipmyn Africa, and SAIMR polyvalent. Surprisingly, the comparison revealed no significant differences between venoms from various African regions and tested antivenoms. It suggests that majority of the toxins present in each venom interacted similarly with tested antivenoms. However, it is possible that there are more subtle differences in venom-antivenom interactions that are difficult to distinguish by quantitative ELISA (Figs 7a, 8a). More insightful information is obtained by plotting the data to compare with each other the protein binding effectiveness of different antivenoms (Fig 7b, 8b). In this case, the differences are evident, and for a dilution of 1:500 in the affinity ELISA test, the highest efficiency in binding venom proteins was demonstrated by EchiTAb+ICP, Antivipmyn Africa and SAIMR polyvalent, respectively (Fig 7b). Recently, Menzies et al. similarly reported higher titre values of EchiTAB+ICP compared to SAIMR polyvalent in the affinity ELISA analysis [17]. Our results also showed higher OD values for Antivipmyn Africa than SAIMR, however, in this case it is essential to note that the experiment was carried out with standardized concentration values, and Antivipmyn Africa was significantly less prediluted than the other two antivenoms. Consequently, a comparison of undiluted products may not present Antivipmyn Africa in an equally favorable manner.

While the comparison of overall venom-antivenom reactivity across different regions did not reveal apparent differences, the data from 2DE Western Blot analysis demonstrated existing variations. Unfortunately, we do not know whether they concern proteins with neurotoxic effects, which stood out as the most distinguishable group of proteins during mass spectrometry analysis. These proteins grouped with other low-molecular-weight toxins, which constitute the majority of the venom, and the gel resolution is insufficient for their clear differentiation. Hence, discernible differences primarily involve proteins constituting a small percentage of the venom, likely having negligible impact on overall toxicity. Examples of such proteins include LAAOs detected in the area A on Eswatini-EchiTAb+ICP membranes, or 5’-nucleotidases, L-amino acid oxidases and metalloproteinases in area F on membranes specific to KwaZulu-Natal venom. Therefore, while these results show variations in both protein composition and efficacy in interacting with different antivenoms, it is challenging to determine their potential impact on overall treatment with antivenom. However, it is probable that the impact might not be substantial due to the low abundance of these proteins in the venom.

Nevertheless, in the context of low-abundance venom proteins, it is necessary to mention those found within region D, which was the most characteristic area differentiating the Limpopo venom from the other two venoms. Despite generating intense signals on Western Blot membranes, these proteins are low in abundance in the venom, as evidenced by the absence of signals on 2D gels stained with Coomassie Brilliant Blue G-250. Their visibility only emerged with silver staining, a more sensitive detection method [21]. This underscores the high antigenicity of these proteins, particularly in the venoms from Eswatini and KwaZulu-Natal. Unfortunately, most signals in this area could not be identified by mass spectrometry despite obtaining promising MS spectra, which is most likely because the sequences of these proteins were absent in the database. The other identified proteins from this region were assigned to three groups: LAAOs, CRISPs, and TNFR (TNF receptor superfamily member 11b). This is unexpected, given that some spots identified as LAAO are around 20 kDa, whereas the smallest reported monomer for LAAO has a mass of 28 kDa [36]. These results, along with those obtained from proteomic venom comparisons, clearly show that incomplete databases are still a major problem in venomics. The recent report by Slagboom et al. clearly illustrated this issue, identifying three times more proteins using species-specific venom gland transcriptome data compared to the UniProt database [37]. This highlights how much data about snake venom composition remains unknown.

To summarize, our study comprehensively analyzed intraspecific differences in *Naja mossambica* venoms from different South African regions, examining their impact on reactivity with commercial antivenoms. Surprisingly, we found a significant presence of neurotoxic proteins in *Naja mossambica* venoms, along with notable variations in composition among different populations. Despite the overall similarity in venom-antivenom reactivity, distinct differences, particularly concerning low-abundance proteins, emerged through 2DE Western Blot analysis, highlighting the intricate nature of venom proteomes.

Limitations of this study should be considered, particularly in the context of venom collection logistics. Venom samples were not directly obtained in Africa, and as such, certain aspects of the variability associated with snake adaptation to their native environments may have been mitigated. However, the consistent collection of venom at various time intervals from snakes within specific regions, coupled with the pooling of samples, served to minimize ontogenic variations, providing a more representative proteomic profile for each population.

## Author Contributions

**Conceptualization:** Konrad K. Hus, Jaroslav Legáth, Aleksandra Bocian

**Data Curation:** Konrad K. Hus

**Formal analysis:** Konrad K. Hus

**Funding acquisition:** Konrad K. Hus, Jaroslav Legáth

**Investigation:** Konrad K. Hus, Justyna Buczkowicz, Monika Pietrowska, Aleksandra Bocian

**Methodology:** Konrad K. Hus, Monika Pietrowska, Aleksandra Bocian

**Resources:** Vladimír Petrilla, Monika Petrillová, Thea Litschka-Koen

**Writing – original draft:** Konrad K. Hus

**Writing – review & editing:** Konrad K. Hus, Justyna Buczkowicz, Monika Pietrowska, Vladimír Petrilla, Monika Petrillová, Jaroslav Legáth, Thea Litschka-Koen, Aleksandra Bocian

## References

[1] World Health Organization. Snakebite envenoming. 2021. Available: https://www.who.int/news-room/fact-sheets/detail/snakebite-envenoming

[2] Gutiérrez JM, Calvete JJ, Habib AG, Harrison RA, Williams DJ, Warrel DA. Snakebite envenoming. Nat Rev Dis Primers 2017;3: 17063. doi: 10.1038/nrdp.2017.79.

[3] Chippaux J-P, Massougbodji A, Habib AG. The WHO strategy for prevention and control of snakebite envenoming: a sub-Saharan Africa plan. J Venom Anim Toxins incl Tropp Dis 2019;25: e20190083. doi: 10.1590/1678-9199-JVATITD-2019-0083.

[4] Farooq H, Bero C, Guilengue Y, Elias C, Massingue Y, Mucopote I, et al. Snakebite incidence in rural sub-Saharan Africa might be severely underestimated. Toxicon 2022;219: 106932. doi: 10.1016/j.toxicon.2022.106932.

[5] Alape-Girón A, Sanz L, Escolano J, Flores-Díaz M, Madrigal M, Sasa M, et al. Snake Venomics of the Lancehead Pitviper *Bothrops asper*: Geographic, Individual, and Ontogenetic Variations. J Proteome Res 2008:7(8): 3556–3571. doi: 10.1021/pr800332p.

[6] Margres MJ, Walls R, Suntravat M, Lucena S, Sánchez EE, Rokyta DR. Functional characterizations of venom phenotypes in the eastern diamondback rattlesnake (*Crotalus adamanteus*) and evidence for expression-driven divergence in toxic activities among populations. Toxicon 2016;119: 28–38. doi: 10.1016/j.toxicon.2016.05.005.

[7] Strickland JL, Smith CF, Mason AJ, Schield DR, Borja M, Castañeda-Gaytán G, et al. Evidence for divergent patterns of local selection driving venom variation in Mojave Rattlesnakes (*Crotalus scutulatus*). Sci Rep 2018;8: 17622. doi: 10.1038/s41598-018-35810-9.

[8] Zancolli G, Calvete JJ, Cardwell MD, Greene HW, Hayes WK, Hegarty MJ, et al. When one phenotype is not enough: divergent evolutionary trajectories govern venom variation in a widespread rattlesnake species. Proc R Soc B 2019;286: 20182735. doi: 10.1098/rspb.2018.2735.

[9] Sousa LF, Holding ML, Del-Rei THM, Rocha MMT, Mourão RHV, Chalkidis HM, et al. Individual Variability in *Bothrops atrox* Snakes Collected from Different Habitats in the Brazilian Amazon: New Findings on Venom Composition and Functionality. Toxins 2021;13(11): 814. doi: 10.3390/toxins13110814.

[10] Rashmi U, Khochare S, Attarde S, Laxme RRS, Suranse V, Martin G, et al. Remarkable intrapopulation venom variability in the monocellate cobra (*Naja kaouthia*) unveils neglected aspects of India’s snakebite problem. J Proteomics 2021;242: 104256. doi: 10.1016/j.jprot.2021.104256.

[11] Smith CF, Nikolakis ZL, Ivey K, Perry BW, Schield DR, Balchan NR et al. Snakes on a plain: biotic and abiotic factors determine venom compositional variation in a wide-ranging generalist rattlesnake. BMC Biol 2023;21: 136. doi: 10.1186/s12915-023-01626-x.

[12] Casewell NR, Jackson TNW, Laustsen AH, Sunagar K. Causes and Consequences of Snake Venom Variation. Trends Pharmacol Sci 2020;41(8): 570–581. doi: 10.1016/j.tips.2020.05.006.

[13] Vermaak SS, Visser A, le Roux TLB. A deadly bed partner: m’Fesi (Mozambique spitting cobra). SA Orthop J 2010;9(4): 58–62.

[14] Petras D, Sanz L, Segura A, Herrera M, Villalta M, Solano D, et al. Snake venomics of African spitting cobras: toxin composition and assessment of congeneric cross-reactivity of the pan-African EchiTAb-Plus-ICP antivenom by antivenomics and neutralization approaches. J Proteome Res 2011;10(3): 1266–80. doi: 10.1021/pr101040f.

[15] Kuna E, Bocian A, Hus KK, Petrilla V, Petrillova M, Legath J, et al. Evaluation of Antifungal Activity of *Naja pallida* and *Naja mossambica* Venoms against Three Candida Species. Toxins 2020;12(8): 500. doi: 10.3390/toxins12080500.

[16] Nguyen GTT, O’Brien C, Wouters Y, Seneci L, Galissà-Calzado A, Campos-Pinto I, et al. GigaScience 2022;11: giac121. doi: 10.1093/gigascience/giac121.

[17] Menzies SK, Litschka-Koen T, Edge RJ, Alsolaiss J, Crittenden E, Hall SR, et al. Two snakebite antivenoms have potential to reduce Eswatini’s dependency upon a single, increasingly unavailable product: Results of preclinical efficacy testing. PLoS Negl Trop Dis 2022;16(9): e0010496. doi: 10.1371/journal.pntd.0010496.

[18] Cox J, Mann M. MaxQuant enables high peptide identification rates, individualized p.p.b.-range mass accuracies and proteome-wide protein quantification. Nat. Biotechnol 2008;26: 1367–1372. doi: 10.1038/nbt.1511.

[19] Tyanova S, Temu T, Sinitcyn P, Carlson A, Hein MY, Geiger T, et al. The Perseus computational platform for comprehensive analysis of (prote)omics data. Nat Methods 2016;13: 731–740. doi: 10.1038/nmeth.3901.

[20] Heberle H, Meirelles GV, da Silva FR, Telles GP, Minghim R. InteractiVenn: a web-based tool for the analysis of sets through Venn diagrams. BMC Bioinformatics 2015;16: 169. doi: 10.1186/s12859-015-0611-3

[21] Shevchenko A, Wilm M, Vorm O, Mann M. Mass spectrometric sequencing of proteins silver-stained polyacrylamide gels. Anal Chem 1996;68(5): 850–858. doi: 10.1021/ac950914h.

[22] Vaudel M, Burkhart J, Zahedi R, Oveland E, Berven FS, Sickmann A, et al. PeptideShaker enables reanalysis of MS-derived proteomics data sets. Nat Biotechnol 2015;33: 22–24. doi: 10.1038/nbt.3109.

[23] Perez-Riverol Y, Bai J, Bandla C, Hewapathirana S, García-Seisdedos D, Kamatchinathan S, et al. The PRIDE database resources in 2022: A Hub for mass spectrometry-based proteomics evidences. Nucleic Acids Res 2022;50(D1): D543–D552. doi: 10.1093/nar/gkab1038.

[24] Tasoulis T, Isbister GK. A current perspective on snake venom composition and constituent protein families. Arch Toxicol 2023;97: 133–153. doi: 10.1007/s00204-022-03420-0.

[25] Bocian A, Ciszkowicz E, Hus KK, Buczkowicz J, Lecka-Szlachta K, Pietrowska M, et al. Antimicrobial Activity of Protein Fraction from *Naja ashei* Venom against *Staphylococcus epidermidis*. Molecules 2020;25(2): 293. doi: 10.3390/molecules25020293.

[26] Hus KK, Marczak Ł, Petrilla V, Petrillová M, Legáth J, Bocian A. Different Research Approaches in Unraveling the Venom Proteome of *Naja ashei*. Biomolecules 2020;10: 1282. doi: 10.3390/biom10091282.

[27] Clare RH, Hall SR, Patel RN, Casewell N. Small Molecule Drug Discovery for Neglected Tropical Snakebite. Trends Pharmacol Sci 2021;42(5): 340–353. doi: 10.1016/j.tips.2021.02.005.

[28] Ljungars A, Laustsen AH. Neutralization capacity of recombinant antivenoms based on monoclonal antibodies and nanobodies. Toxicon 2022;222: 106991. doi: 10.1016/j.toxicon.2022.106991.

[29] Menzies SK, Arinto-Garcia R, Amorim FG, Cardoso IA, Abada C, Crasset T, et al. ADDovenom: Thermostable Protein-Based ADDomer Nanoparticles as New Therapeutics for Snakebite Envenoming. Preprints: 2023101070 [Preprint]. 2023 [cited: 2023 October 19]. Available from: 10.20944/preprints202310.1070.v1

[30] Potet J, Smith J, McIver L. Reviewing evidence of the clinical effectiveness of commercially available antivenoms in sub-Saharan Africa identifies the need for a multi-centre, multi-antivenom clinical trial. PLOS Negl Trop Dis 2019;13(6): e0007551. doi: 10.1371/journal.pntd.0007551.

[31] Amorim FG, Redureau D, Crasset T, Freuville L, Baiwir D, Mazzucchelli G, et al. Next-Generation Sequencing for Venomics Application of Multi-Enzymatic Limited Digestion for Inventorying the Snake Venom Arsenal. Toxins 2023:15(6): 357. doi: 10.3390/toxins15060357.

[32] Khan NA, Amorim FG, Dunbar JP, Leonard D, Redureau D, Quinton L, et al. Inhibition of bacterial biofilms by the snake venom proteome. Biotechnol. Rep 2023;39: e00810. doi: 10.1016/j.btre.2023.e00810.

[33] Galea CA, Nguyen HM, Chandy GK, Smith BJ, Raymond RS. Domain structure and function of matrix metalloprotease 23 (MMP23): role in potassium channel trafficking. Cell Mol Life Sci 2014;71: 1191–1210. doi: 10.1007/s00018-013-1431-0.

[34] Zhang XW, Wang Y, Wang XW, Wang L, Mu Y, Wang JX. A C-type lectin with an immunoglobulin-like domain promotes phagocytosis of hemocytes in crayfish *Procambarus clarkii*. Sci Rep 2016;6: 29924. doi: 10.1038/srep29924.

[35] Zhou K, Qin Y, Song Y, Zhao K, Pan W, Nan X, et al. A Novel Ig Domain-Containing C-Type Lectin Triggers the Intestine-Hemocyte Axis to Regulate Antibacterial Immunity in Crab. J Immunol 2022;208(10): 2343–2362. doi: 10.4049/jimmunol.2101027.

[36] Zuliani JP, Paloschi MV, Pontes AS, Boeno CN, Lopes JA, Setubal SS, et al. Reptile Venom L-Amino Acid Oxidases – Structure and Function. In: Mackessy SP, editor. Handbook of Venoms and Toxins of Reptiles. Second Edition. CRC Press; 2021. pp. 413–430.

[37] Slagboom J, Derks RJE, Sadighi R, Somsen GW, Ulens C, Casewell NR, et al. High-Throughput Venomics. J Proteome Res. 2023;22(6): 1734–1746. doi: 10.1021/acs.jproteome.2c00780.

